# Developmentally Regulated Genome Editing in Terminally Differentiated N_2_-Fixing Heterocysts of *Anabaena cylindrica* ATCC 29414

**DOI:** 10.1101/629832

**Authors:** Yeyan Qiu, Liping Gu, Shengni Tian, Jagdeep Sidhu, Jaimie Gibbons, Trevor Van Den Top, Jose L. Gonzalez-Hernandez, Ruanbao Zhou

**Author notes:** Correspondences: Jose L. Gonzalez-Hernandez, Ruanbao Zhou.

## Abstract

Some vegetative cells of *Anabaena cylindrica* are programed to differentiate semi-regularly spaced, single heterocysts along filaments. Since heterocysts are terminally differentiated non-dividing cells, with the sole known function for solar-powered N_2_-fixation, is it necessary for a heterocyst to retain the entire genome (≈7.1 Mbp) from its progenitor vegetative cell? By sequencing the heterocyst genome, we discovered and confirmed that at least six DNA elements (≈0.12 Mbp) are deleted during heterocyst development. The six-element deletions led to the restoration of five genes (*nifH1*, *nifD*, *hupL*, *primase P4* and a hypothetical protein gene) that were interrupted in vegetative cells. The deleted elements contained 172 genes present in the genome of vegetative cells. By sequence alignments of intact *nif* genes (*nifH*, *nifD* and *hupL*) from N_2_-fixing cyanobacteria (multicellular and unicellular) as well as other N_2_-fixing bacteria (non-cyanobacteria), we found that interrupted *nif* genes all contain the conserved core sequences that may be required for phage DNA insertion. Here, we discuss the *nif* genes interruption which uniquely occurs in heterocyst-forming cyanobacteria. To our best knowledge, this is first time to sequence the genome of heterocyst, a specially differentiated oxic N_2_-fixing cell. This research demonstrated that (1) different genomes may occur in distinct cell types in a multicellular bacterium; and (2) genome editing is coupled to cellular differentiation and/or cellular function in a heterocyst-forming cyanobacterium.

## Introduction

Nitrogen is one of the most important, abundant life elements in bio-macromolecules such as DNA, RNA, and proteins. Although nearly 80% of air is N_2_ gas, most of organisms are unable to use this form of dinitrogen (N_2_) to make these essential bio-macromolecules. Fortunately, some cyanobacteria can photosynthetically fix atmospheric N_2_ gas into a form (ammonia) that can be used by other organisms. Through billions of years of evolution, *Anabaena* species has gained the unique capability of using solar energy to reduce atmospheric N_2_ to ammonia in specially differentiated N_2_-fixing cells called heterocysts^1, 2^. The sole function for heterocysts is its solar-powered, oxic N_2_-fixation. Thus, heterocysts also offer scientists a rare opportunity to unlock the mystery of genome requirements for photosynthetic N_2_-fixation. Unlocking genomic secrets of heterocysts would help guide scientists to genetically engineer crops (leaves) to make self-fertilizing plants/crops using sunlight and atmospheric N_2_ gas, just as heterocysts have done for billions of years.

Regardless of nitrate availability in its growth medium, some vegetative cells of *Anabaena cylindrica* ATCC 29414 (hereafter *A. cylindrica*) can initiate a development program to form heterocysts that are present singly at semi-regular intervals along the filaments ^3^. By sequestering nitrogenase within heterocysts, *A. cylindrica* can carry out the two incompatible biochemical processes simultaneously: O_2_-producing photosynthesis and O_2_-labile N_2_ fixation. Heterocyst-based N_2_-fixation is a uniquely oxic, solar-powered process, which is distinct from anaerobic N_2_-fixation present in other bacterial species. This provides great potential for application in agriculture compared to all other N_2_-fixing bacteria, which are unable to use solar energy and also require anaerobic conditions.

Unlike vegetative cells, heterocysts are terminally differentiated cells that have two extra O_2_-impermeable layers of glycolipids and polysaccharides to exclude the O_2_, which inactivates nitrogenase ^4, 5^. Heterocysts are morphologically and biochemically specialized for solar-powered, oxic N_2_-fixation. The heterocysts normally develop and mature within 24 hours. This 24-h period during which heterosysts are differentiating from vegetative cells is called the pro-heterocyst stage. A mature heterocyst is larger, more rectangular in shape with less granular cytoplasm than a vegetative cell, and it has thickened cell walls and a refractive polar granule at each end of the cell ^6^. Cells with these characteristics, but lacking the thickened cell walls and the polar granules, are counted as pro-heterocysts ^7^.

Many genes have been identified to be involved in regulating heterocyst differentiation. HetR is a master transcription regulator specifically required for heterocyst differentiation ^8, 9, 10^. Several other regulatory genes such as *nrrA* ^11^, *ccbP* ^12^, *hetN* ^13^, *hetF*, *patA* ^14^, *patN* ^15^, *patU* ^16^, *hetZ* ^17^, *patS* ^18, 19^, *hepK* ^5^, and *hetP* ^20^ were also found to play critical roles during heterocyst differentiation. During heterocyst development in *Anabaena* sp. PCC 7120, at least three DNA elements (11-kb, 55-kb and 9.4-kb) inserted within *nifD*, *fdxN* and *hupL*, respectively, are programmed to excise from the heterocyst genome by developmentally regulated site-specific recombination ^21, 22, 23^. Both deletions of the 11-kb and 55-kb elements had been proven to be necessary for the heterocyst-based N_2_-fixation, but not required for the differentiation of heterocysts in *Anabaena* sp. PCC 7120^23, 24^, while the 9.4-kb deletion effects neither N_2_-fixation nor heterocyst formation^25^.

The nitrogenase complex is encoded by a group of genes called *nif* genes. Many heterocyst-forming cyanobacteria have *nif* (*ni*trogen *f*ixation) genes (e.g., *nifH*, *nifD*, *nifK*, *fdxN*) interrupted by DNA elements that must be excised during heterocyst development ^21, 26^. Since heterocysts are terminally differentiated, non-dividing cells, with the sole function of solar-powered N_2_-fixation, is it necessary for a heterocyst to retain the entire genome (≈7.1 Mbp) from its progenitor vegetative cell? To answer this question, we isolated heterocysts from *A. cylindrica* and sequenced the genomic DNA from heterocysts and vegetative cells. After mapping the NGS-based 286 heterocyst contigs of *A. cylindrica* ATCC 29414 to the reference genome of *A. cylindrica* PCC 7122 NCBI (GCA_000317695.1) using BLAST-N (E-value < 1 ×10^−150^), six DNA elements (≈0.12 Mbp) were found to be deleted from the heterocyst genome during heterocyst development. The six-element deletions in heterocysts led to a loss of 172 genes, but restored five genes (*nifH1*, *nifD*, *hupL*, *primase P4*, a hypothetical protein gene) that were interrupted in vegetative cells.

## Methods

### Isolation and purification of heterocysts

Heterocysts were obtained from *A. cylindrica* grown in 5 L AA/8 medium free of combined nitrogen ^27^, shaking at 150 rpm under illumination (50-60 μE·m^−2^·s^−1^ at the culture surface) for 7 days to an OD_700_ of 0.03. Cultures were harvested by centrifugation at 6,000 x g for 15 min and re-suspended in 80 mL ddH_2_O. Vegetative cells were disrupted by passing suspensions through a Nano DeBEE 30 High Pressure Homogenizer (BEE International) at 15,000 psi (lb/in^2^) three times. The suspensions were centrifuged down at 4,000 x g for 10 min. To separate the debris of vegetative cells from heterocysts, pellets were re-suspended in 1 mL ddH_2_O, and the suspension was centrifuged at 1,100 x g for 5 min. Two layers were formed: a bottom green pellet and a top loose yellow pellet. The top yellow pellet of vegetative cells debris was discarded and the bottom green heterocysts pellet was washed by re-suspending with 1 mL ddH_2_O and re-centrifuging at 1,100 x g for 5 min. This wash step was repeated 4 times. After each washing step, the heterocyst fraction was checked microscopically in order to ensure that heterocysts were pure. The purified heterocysts were stored at −80°C.

### Isolation of genomic DNA

*Anabaena cylindrica* ATCC 29414 was grown in 50 mL AA/8 medium (free of combined nitrogen) and AA/8N (nitrate-containing medium) ^27^ for 7 days, and OD_700_ were 0.03 and 0.028, respectively. To extract the DNA from vegetative cells, the cultures were centrifuged at 13,000 x g for l5 min; 500 μL of 10% sucrose buffer (50 mM Tris-HCl, pH 8.0, 10 mM EDTA) was used to suspend the cell pellets; 50 μL of 125 mg/mL lysozyme (Sigma), 150 μL 10% SDS and 10 μL RNase of 10 mg/mL were added. The ≈800 μL suspension was incubated at 37°C for 1 hr. Then the total amount of the suspension was measured, an equal amount of saturated phenol (pH 6.6±0.2) was added, and the reagents were mixed by vortexing. The suspension was centrifuged at 13,000 x g at room temperature for 10 min. The top aqueous solution was transferred to a new 1.5 mL Eppendorf tube and an equal amount of chloroform solution (chloroform: isoamyl aclcohol = 24:1) was added. The tube was vortexed and the suspension was centrifuged at 13,000 x g at room temperature for 10 min. The top aqueous solution was then transferred to new tube and an equal volume of pre-cold isopropanol was added to precipitate total DNA. Some white pellet was obtained after centrifuging at 13,000 x g at 4°C for 10 min. The supernatant was discarded, and the pellet was washed first with 70% ethanol and then 95% ethanol. The white pellet was air-dried for 5 min, and a final 30 μL of ddH_2_O was added to dissolve the total DNA.

To break the heterocysts and extract its genomic DNA, the purified 13.1 mg (wet weight) of heterocysts stored at −80°C were re-suspended in 500 μL 10% sucrose buffer (50 mM Tris-HCl, pH 8.0, 10 mM EDTA). The suspension was centrifuged at 13,000 x g for 5 min. After removing the supernatant, another 500 μL 10% sucrose buffer was added to suspend the pellets, and 50 μL of 125 mg/mL lysozyme was added. The total suspension was incubated at 37°C for 1.5 hr. The suspension was sonicated at 70% amplitude for 10 s with 0.5 s pulse on and 0.5 s pulse off. The sonication process was repeated 3 times. TissueLyser II (Qiagen) was further used to break heterocysts at frequency of 30/s for 8 min. The sample was frozen in liquid nitrogen immediately for 2 min and defrosted at 80°C for 2 min. The TissueLyser, liquid nitrogen freezing, defreezing processes were repeated for another 3 times. Then 150 μL of 10% SDS, 150 μL of 0.5 M EDTA (pH 8.0) were added to the suspension and incubated at 80°C for 30 min. Ten μL of RNase (10 mg/mL) was added into the tube and incubated at room temperature for 15 min. Next, the heterocyst DNA extraction procedures followed the same saturated-phenol method as described for vegetative cells. A final volume of 15 μL of ddH_2_O was used to dissolve heterocyst DNA. All the DNA was quantified by Qubit 3.0 (Thermo scientific).

### Genome sequencing

Sequencing libraries for the vegetative and heterocyst DNA samples were produced using a Nextera XT library preparation kit (Illumina) following the protocol described by the manufacturer. Libraries were quantified using a Qubit 3.0 and quality checked with an Agilent Bioanalyzer (Agilent). Equimolar amounts of both libraries were loaded as part of an Illumina NextSeq 500 high output run producing 2×150 bp paired ends reads. Libraries preparation and sequencing was done at the South Dakota State University Genomics Sequencing Facility.

### Bioinformatics analysis

Reads trimming, assembly and mapping were carried out using CLC Genomics Workbench 10.1.1 (Qiagen). Trimming was carried out using a Q of 20 as the cutoff, eliminating any read with any ambiguous nucleotide and removing the 5 and 15 terminal nucleotides in the 3’ and 5’ ends respectively. Assembly of the trimmed reads was accomplished by setting an arbitrary minimum contig length of 3,000 bp and using the automated function to select a word size of 23 and a bubble size of 50; finally, reads were mapped (mismatch cost: 2, insertion cost: 3, deletion cost: 3, minimum length fraction: 0.5 and minimum similarity fraction: 0.9) to the assembly and the results used to correct the contigs sequences. To detect possible deletions, trimmed reads were mapped to the reference genome of *A. cylindrica* PCC 7122 using the large gap read mapper function (mismatch cost: 2, insertion cost: 3, deletion cost: 3, minimum length fraction: 0.9, minimum similarity fraction: 0.95 and randomly assigning those reads mapping in multiple locations).

To determine the phylogenetic distance of our *A. cylindrica* ATCC 29414 and reference genome, both contigs from the vegetative cells and heterocysts assemblies were compared to the four copies of 16S rRNA gene sequences in *A. cylindrica* PCC 7122 (GCA_000317695.1). The four copies are *Anacy_R0013*, *Anacy_R0015*, *Anacy_R0054* and *Anacy_R0070*. With the high fidelity of these two genomes, we compared our assemblies of vegetative cells and heterocysts to *A. cylindrica* PCC 7122 with a cutoff E-value 1E-150 through Linux command line version of BLAST+. Granges^28^ was further used to discover the unique predicted deletions in heterocysts. The genes found in these deletion regions were annotated with *A. cylindrica* PCC 7122 reference genome.

### PCR confirming the edited genes

Specific primers ZR1676 (0.5 μM) and ZR1677 (0.5 μM) were used to amplify intact *nifH1* using genomic DNA from heterocysts and vegetative cells of *A. cylindrica* ATCC 29414. The 891bp band was extracted using a DNA extraction kit (Qiagen), and cloned into pCR2.1-TOPO vector (Invitrogen). The colony PCR confirmed the correct clones (plasmids) were extracted and sent for DNA sequencing. The intact *nifD*, *hupL*, *primase P4* and a hypothetical gene of joined *anacy_RS29550* and *anacy_RS29775* were PCR amplified by Phusion High-Fidelity DNA Polymerase (NEB) with specific primers listed in Table S1. The PCR products amplified with genomic DNA from heterocysts and vegetative cells of *A. cylindrica* were purified with the Qiagen PCR clean kit for DNA sequencing.

### Quantitative polymerase chain reaction (qPCR)

Quantitative PCR was performed to determine the ratios of the edited genome vs unedited genome in heterocysts. For qPCR, vegetative cells were grown in AA/8 or AA/8N, and two pairs of primers were used for each individual gene (primers listed in Table S1). Ten ng DNA isolated from vegetative cells grown in AA/8 and AA/8N and 10 ng DNA isolated from heterocysts were added to a 20-μL reaction containing 0.2 units of Phusion High-Fidelity DNA Polymerase (NEB), 1X Phusion buffer, dNTP (0.25 mM), and primers (0.5 μM). Each qPCR reaction had 5 replicates. The qPCR program was: 95°C for 10 min; 40 cycles of 95°C for 30 s, 55°C for 30 s, 72°C for 30 s; and a dissociation stage of 95°C for 15 s, 55°C for 30 s, 95°C for 15 s.

### Determining the frequency of heterocysts

*A. cylindrica* can form heterocysts in both AA/8 (without combined nitrogen) and AA/8N (with combined nitrogen). The same cultures used for isolation of genomic DNA (above) were used to determine the frequency of heterocysts using microscopy accounting. Pictures were taken and the total numbers of heteocysts and vegetative cells were counted to determine the heterocyst frequency in both AA/8 and AA/8N growth media ^27^.

## Results

### Isolation and purification of heterocysts for genome sequencing

Approximately 4.46% of *A. cylindrica* vegetative cells grown in AA/8 medium (free of combined nitrogen) ^27^ can form single heterocysts (Fig. S1a), and the heterocysts were purified to a purity of 99.52 ± 0.48% (Fig. S1b). Unlike the other heterocyst-forming cyanobacteria, such as *Anabaena* sp. PCC 7120, *Anabaena variablis* ATCC 29413 and *Nostoc punctiforme* ATCC 29133, *A. cylindrica* can also form single heterocysts with a frequency of ≈2.04% (Fig. S1c) when grown in a nitrate-containing medium, such as AA/8N ^27^.

### Comparative genomic sequencing between vegetative cells and heterocysts identified DNA deletions in heterocysts genome

#### NGS of genomic DNA from vegetative cells and heterocysts

Although two *A. cylindrica* genomic sequence databases are available, which are the complete genome of *A. cylindrica* PCC 7122 in NCBI (GCA_000317695.1) and the 154 contigs of *A. cylindrica* ATCC 29414 from Dr. John C. Meeks at UC Davis (http://scorpius.ucdavis.edu/gmod/cgi-bin/site/anabaena02?page=assembly), there is no report for heterocyst genomic sequencing data. Here, we sequenced the genomic DNA isolated from highly purified heterocysts (Fig. S1b) and vegetative cells, respectively. The complete genome of *A. cylindrica* PCC 7122 was used as the reference genome for short-read mapping (Fig. 1a & Fig. S4) and contig assembling for the heterocyst genome and vegetative cell genome of *A. cylindrica* ATCC 29414. A total of 254 contigs (VC254) from vegetative cells and 286 contigs (HT286) from heterocysts were assembled by CLC genomic workbench 11 (Qiagen). The total accumulative length of VC254 contigs is 6,761,576 bp with N50 for 43,156 bp, where the largest contig is 177,696 bp. The total accumulative length of HT286 contigs is 6,756,227 bp with N50 is 37,602 bp, where the largest contig is 181,707 bp.

**Fig. 1.**
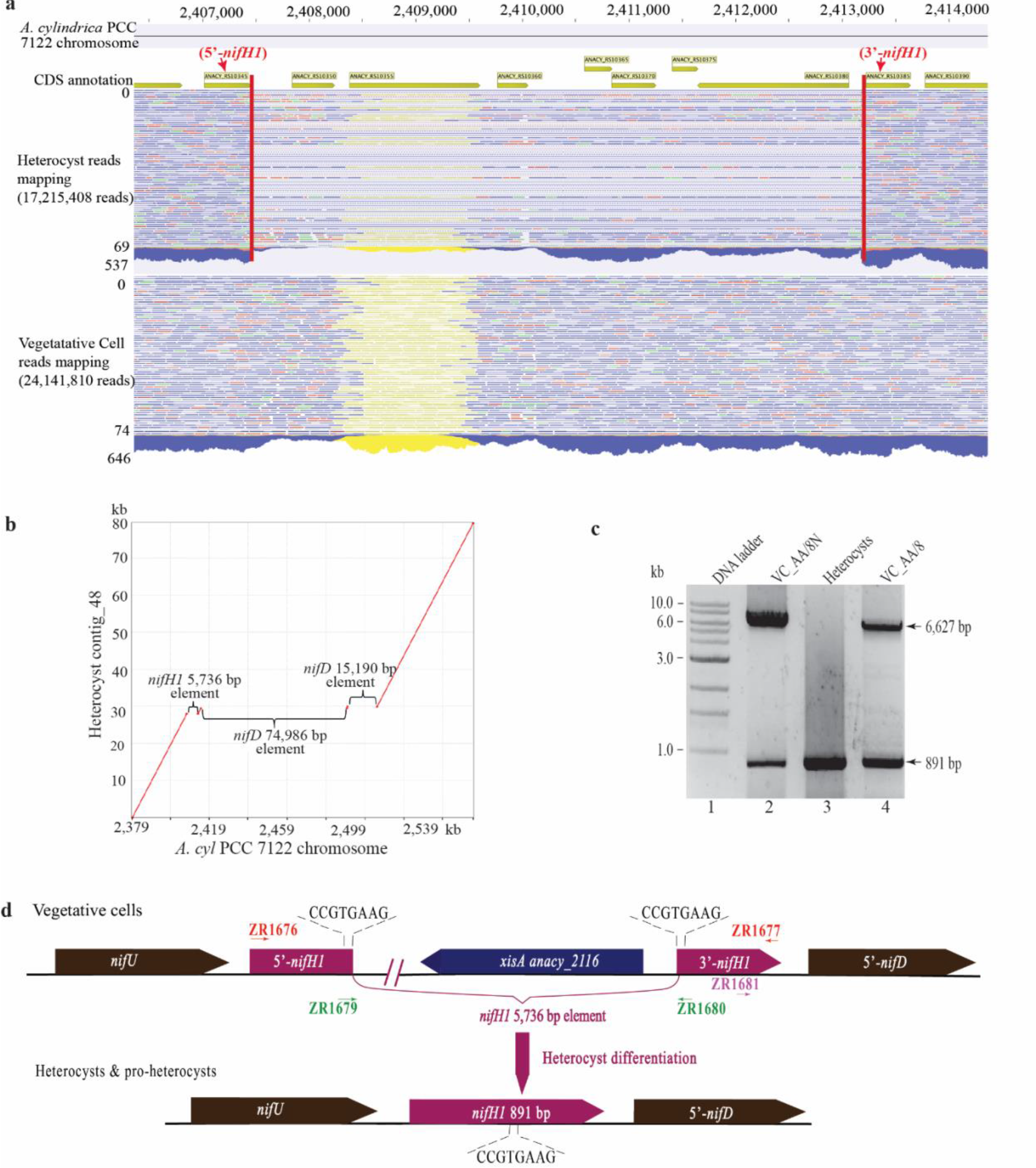
The *nifH1* 5,736 bp element deletion in *A. cyl* ATCC 29414. **a)** The 150 bp paired reads mapping of heterocyst and vegetative cells to reference genome *A. cylindrica* PCC 7122. The blue lines represent the paired reads, the red lines represent the unpaired forward reads, the green lines represent the unpaired reverse reads and the yellow lines represent the unspecific mapping reads. The region between vertical red lines in the heterocyst mapping indicates the 5,736 bp deletion. The sequencing depths of this region were 537 and 646 short reads, respectively, for heterocyst and vegetative cells. In this figure, 69 of 537 and 74 of 646 are shown. The dash lines in this region show that these reads mapping to the *A. cylindrica* PCC 7122 genome were interrupted. In other words, this 5,736 bp *nifH1*-element was confirmed to be deleted with these short reads from heterocysts. *Anacy_RS 10345* and *Anacy_RS 10385* are the truncated 5’-*nifH1* and 3’-*nifH1*, respectively. **b)** The MUMmer plot^29^ mapping of heterocyst contig_48 to the *A. cylindrica* PCC 7122 genome (region of 2,379 kb to 2,555 kb were shown). The missing regions of 5,736 bp *nifH1*-elemnt, 74,986 bp 5’-*nifD*-elemnt and 15,190 bp 3’-*nifD*-elemnt in the plot mapping confirmed these deletions were observed in the heterocyst contig_48. **c)** The PCR products amplified with primers ZR1676/ZR1677 and DNA extracted from vegetative cells grown in AA/8N (with fixed nitrogen), AA/8 (free of fixed nitrogen) and heterocysts. Two bands of 6,627 bp and 891 bp were obtained in both vegetative cells, while in heterocysts only the 891 bp band was present. **d)** The schematic picture of *nifH1* element deletion. The 5,736 bp *nifH1*-element between 5’-*nifH1* and 3’-*nifH1* was removed during heterocyst differentiation. The restored intact *nifH1* was present in heterocysts and perhaps in pro-heterocysts. The integrase XisA’s (Anacy_2116) recognition core sequence (the direct repeat sequence flanking the DNA element) is CCGTGAAG; this sequence may be required for specific phage DNA insertion.

By BLASTing the VC254 contigs against the PCC 7122 genome with cutoff E-value 1E-150, our VC254 contigs produced 6,819,509 bp uniquely matching the PCC 7122 genome (7,063,285 bp), a 96.55% coverage of PCC 7122 genome sequence. Among the 6,819,509 bp of ATCC 29414 VC254 contigs, subtracting the mismatched bps and gap-open bps in the original BLAST data (Table S2), we found that the ATCC 29414 genome sequence is nearly identical (at least 99.63% identity) to the PCC 7122 complete genome. Similarly, our HT286 contigs produced 6,787,777 bp, a 99.1% coverage of the PCC 7122 genome sequence with at least a 99.60% identity (Table S3). The four copies of the 16S rRNA gene sequence of ATCC 29414 were, 99.723%, 99.797%, 99.932%, 100%, respectively, identical to those of PCC 7122 (Table S4). Based on our comparative genomics analysis between two *A. cylindrica* strains, we conclude that these two strains are nearly identical genetically.

#### Identification of six DNA element deletions in heterocysts by sequencing heterocyst genome and PCR confirmation

Although we are currently unable to assemble both vegetative cells and heterocysts of *A. cylindrica* ATCC 29414 into single pseudomolecules, we discovered six major deletions in the heterocyst genome using both contigs mapping and BLAST-N. The six major DNA element deletions were identified in heterocyst contigs (Table 1, group I contigs). They are Δ5,736 bp in *nifH*1 (Fig. 1a-d, Table 1), Δ74,986 bp in 5’-*nifD* and Δ15,190 bp in 3’-*nifD* (Fig. 1b, Fig. 3a,e & Table 1), Δ59,225 bp in primase P4 (Fig. 3b,e & Table 1), Δ20,842 bp in *hupL* (Fig. 3c,f & Table 1), Δ39,998 bp in hypothetical gene of *pANACY. 03* (Fig. S3d, g & Table 1). Consistent with the short-read mapping results (Fig. 1a & Fig. S4), the heterocysts contigs assembly also identified two populations (groups I & II) of contigs (Table 1). The contigs in group I confirmed these six DNA element deletions and the contigs in Group II (Table 1) remained unedited (i.e., contained these six DNA elements identified in the vegetative contigs) (Table 1). These two groups of contigs may indicate the heterogeneity of the heterocyst genome. The 8 contig sequences in group I (with deletions of the above six DNA elements), containing the restored five intact genes (Table 1) and their flanking sequences, have been deposited to GenBank. Their access Nos. are MG594022 through MG594031 (Table S5). The MUMmer mapping ^29^ of Group I contigs against the reference genome (*A. cylindrica* PCC 7122) were also performed and are shown in Fig. 1b and Fig. S2.

**Table 1.** *Anabaena cylindrica* ATCC 29414 VC sequence contigs and HT sequence contigs coverage of the six DNA elements compared to the complete genome of *A. cylindrica* PCC 7122

| Six DNA elements | Element length (bp) | Vegetative cells (VC) contigs |  | Heterocyst (HT) contigs |  |  |
| --- | --- | --- | --- | --- | --- | --- |
|  |  | coverage (%) | identity (%) | Group I coverage (%) | Group II coverage (%) | Group II identity (%) |
| *hupL* | 20,842 | 20,842 bp (100%) | 99.996% | contig_130&339 (0) | 19,200bp (92%) | 99.995% |
| 3'-*nifD* | 15,190 | 15,190 bp (100%) | 97.384% | contig_53 (0) | 0 bp (0%) | 0 |
| 5'-*nifD* | 74,986 | 70,921bp (95%) | 99.315% | contig_48&83 (0) | 71,564bp (95%) | 99.301% |
| *nifH1* | 5,736 | 5,121bp (90%) | 99.481% | contig_48 (0) | 4,979bp (87%) | 99.508% |
| Primase P4 | 59,225 | 55,323bp (93%) | 99.138% | contig_288&303 (0) | 55,973bp (95%) | 99.112% |
| hypothetical protein | 39,998 | 39,639bp (99%) | 100% | contig_30 (0) | 39,990bp (99%) | 99.990% |

After mapping 286 heterocyst contigs of ATCC 29414 to the reference chromosome (6,395,836 bp) of PCC 7122 (NC_019771.1), we further confirmed the five chromosomal deletions (totalized 116,754 bp) of large DNA elements (*hupL*, *nifH1*, *primase P4*, *5’-nifD* and *3’*-*nifD*) in the heterocyst chromosome. In addition, there were also 33 bigger gaps (gap sizes vary from 500 bp to 6,450bp) or potential deletions (PD) mapped to the reference chromosome of PCC 7122 (Fig. S3 & Table S6a).

### Quantitative PCR confirmed restoration of five intact genes in heterocyst genome

#### nifH1 restoration editing in heterocysts and vegetative cells

Two PCR products (6,627 bp and 891 bp) were amplified with genomic DNA from vegetative cells grown in AA/8N (lane 2 in Fig. 1c) and AA/8 (lane 4) with primers ZR1676, ZR1677 (Table S1), while only one product of 891 bp was amplified in heterocyst genomic DNA (lane 3 in Fig. 1c). These three 891 bp PCR product sequences were confirmed to be identical to the coding region of *nifH1* in the heterocyst contig_48 (GenBank accession #: MG594028). In other words, both the heterocyst genomic sequence contig_48 and the intact *nifH1* PCR product sequence confirmed that a 5,736 bp element disrupts the *nifH1* gene in vegetative cells, but is precisely excised to restore the intact *nifH1* in heterocysts. This 5,736 bp element contains the direct repeat sequence CCGTGAAG at both ends inserted within *nifH1*. Interestingly, the intact *nifH1* was also amplified from the genomic DNA isolated from vegetative cells grown in AA/8N and AA/8 (Fig. 1c, lanes 2 & 4). Further quantitative PCR (qPCR) with specific primers (Table S1) targeting the edited *nifH1* and total *nifH1* (both unedited and edited) was performed with genomic DNA isolated from different types of cells. The qPCR data (Fig. 2, *nifH1*) showed that 100±0.77% of interrupted *nifH1* was edited to be intact in genome of heterocysts, while only 11.92±0.86% (AA/8) and 2.89±0.26% (AA/8N) of interrupted *nifH1* was edited to be intact in genome of vegetative cells grown in the medium without combined nitrogen (AA/8) and with combined nitrogen (AA/8N).

**Fig. 2.**
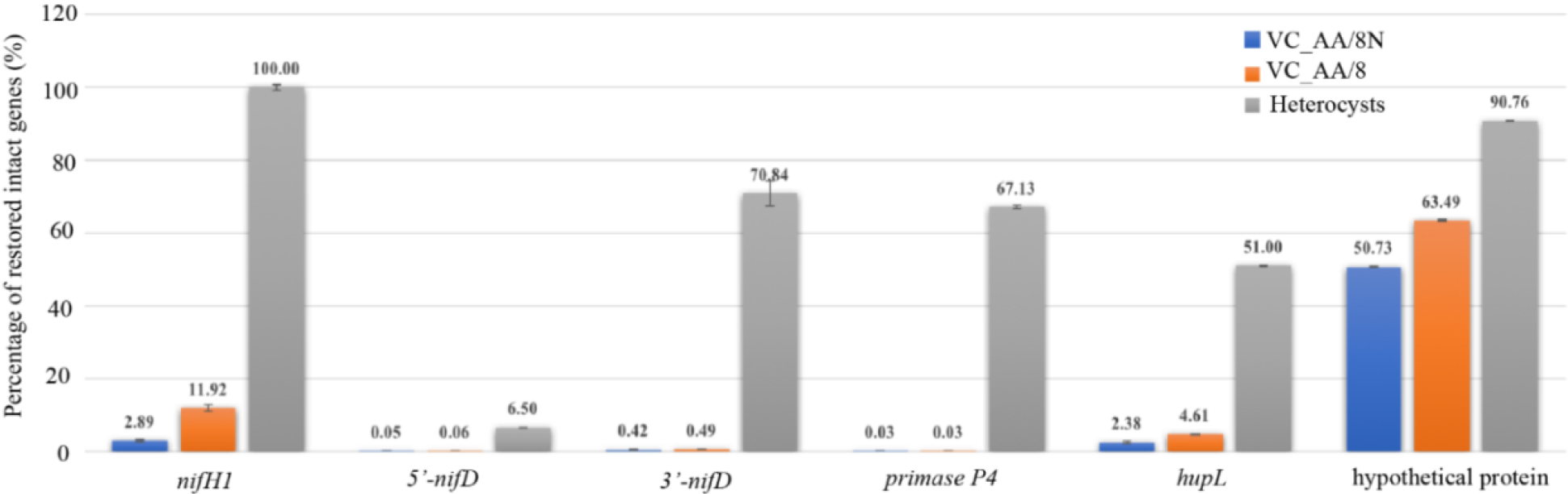
The percentages of restored intact genes of *nifH1*, 5’-*nifD*, 3’-*nifD*, *primase P4*, *hupL* and a hypothetical protein gene joined by *anacy_RS29550* & *RS_29775* in vegetative cells (grown in AA/8N, AA/8) and heterocysts. The total of each gene (interrupted and intact), and the intact only were determined by qPCR using two sets of primers (seen table S1. *nifH1*, total: ZR1681&ZR1677, edited intact *nifH1*: ZR1679&ZR1680; *nifD*, total: ZR1735&ZR1737, 5’-*nifD* element deletion: ZR1738&1739 (Fig. 3), 3’-*nifD* element deletion: ZR1740&ZR1736 (Fig. 3); *primase P4*, total: ZR1749&ZR1750 (Fig. 3), edited intact *primase P4*: ZR1751&ZR1752 (Fig. 3); *hupL*, total: ZR1743&ZR1734, (Fig.3) edited intact *hupL*: ZR1741&ZR1742 (Fig.3); jointed *anacy_RS29550* & *RS_29775* hypothetical protein gene, total: ZR1744&ZR1747 (Fig.3); edited intact: ZR1745&ZR1746 (Fig. 3). The percentages of each restored intact gene were calculated by 2^^[-(Ct(intact)-Ct(total)].^ Except for the jointed *anacy_RS29550* & *RS_29775* hypothetical protein gene, the percentages of intact gene for other 5 genes were significantly lower in vegetative cells than in heterocysts. The editing ratios in vegetative cells grown in AA/8 media were generally higher than in AA/8N.

#### nifD restoration editing in heterocysts and vegetative cells

A 1,544 bp fragment was amplified with genomic DNA (Fig. S2a) from heterocysts (lane 4) as well as from vegetative cells grown in AA/8 (lane 3) or AA/8N (lane 2) with primers ZR1735, ZR1736. These three PCR products of *nifD* were sequenced and confirmed to be identical to the coding region of intact *nifD* (GenBank accession #: MG594028). In other words, two DNA elements (74,986 bp 5’-*nifD* element, 15,190 bp 3’-*nifD*-element) inserted within *nifD* were precisely removed to restore an intact *nifD* (1,503 bp) (Top panel in Fig. 3e). The qPCR data (Fig. 2) showed that 6.50±0.01% of 5’-*nifD* was edited (a 74,986 bp 5’-*nifD* element was precisely removed) in heterocysts, while the edited 5’-*nifD* in vegetative cells accounted for only 0.06±0.01% (AA/8) and 0.05%±0.01 (AA/8N) of total *nifD*. For 3’-*nifD* editing, 70.84±3.4% of 3’-*nifD* was edited (a 15,190 bp 3’-*nifD*-element was precisely removed) in heterocysts, while in vegetative cells the edited 3’-*nifD* accounted for only 0.49±0.03% (AA/8) and 0.42±0.05% (AA/8N).

**Fig. 3.**
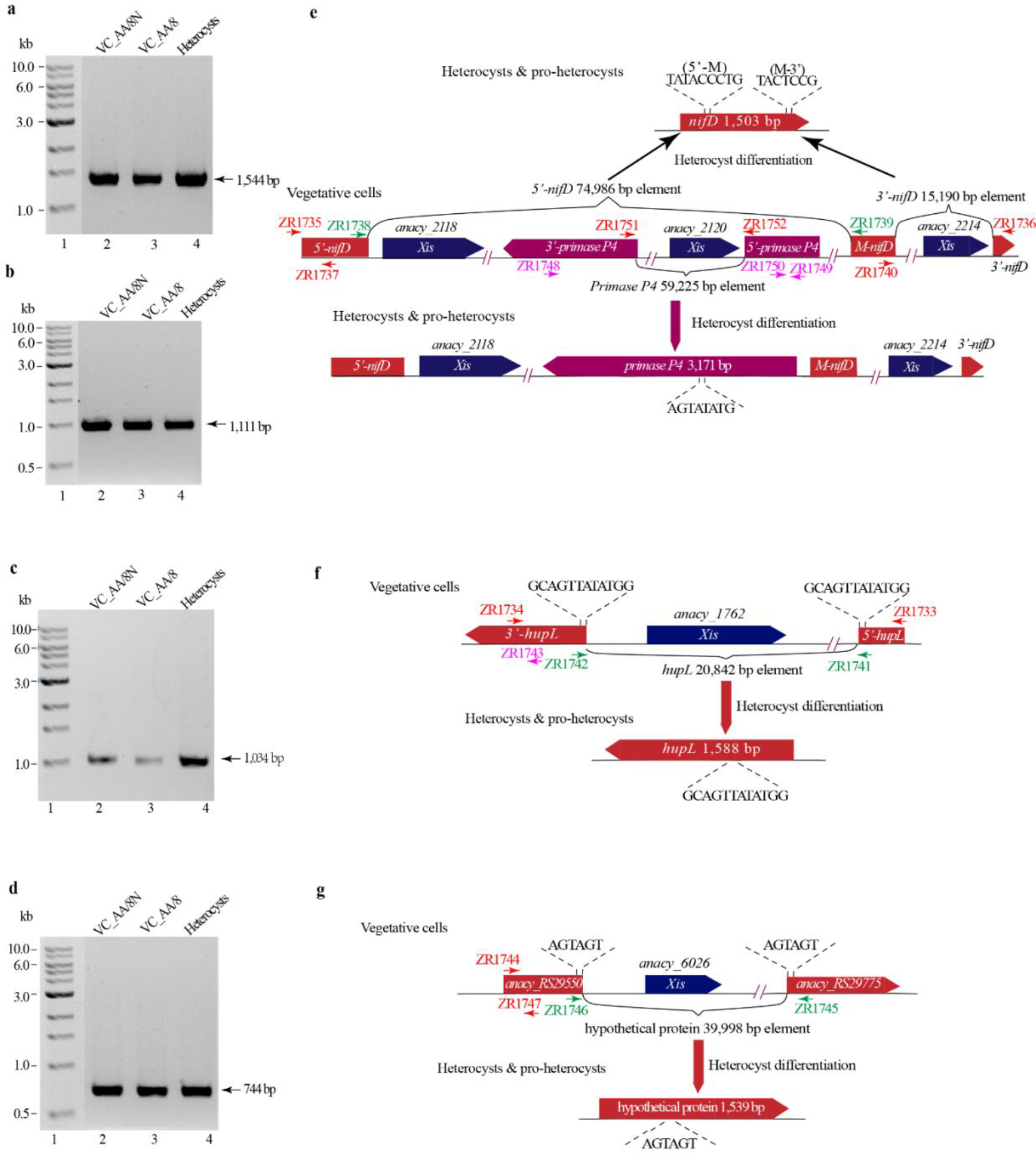
The PCR confirmation of the deletion in *nifD*-, *primase P4*-, *hupL*-, jointed *anacy_RS29550* & *RS_29775* hypothetical protein-DNA elements (a-d) and the deletion scheme were shown (e-g). **a)** A 1,544 bp band was amplified in both vegetative cells and heterocysts DNA with primers ZR1735/ZR1736, which confirmed the 3 *nifD* elements’ deletion were observed in some vegetative cells and heterocys. **b)** A 1,111 bp band was amplified in both vegetative cells and heterocysts DNA with primers ZR1748/ZR1749, which confirmed the *primase P4* element deletion were observed in some vegetative cells and heterocysts. **c)** A 1,034 bp band was amplified in both vegetative cells and heterocysts DNA with primers ZR1733/ZR1734, which confirmed the *hupL* element deletion. **d)** A 744 bp band was amplified in both vegetative cells and heterocysts DNA with primers ZR1744/ZR1745, which confirmed the jointed *anacy_RS29550* & *RS_29775* hypothetical protein 39,998 bp element deletion in *pANACY. 03*. **e)** The upper of this scheme showed the formation of intact *nifD* by removing the 5’-*nifD* and 3’-*nifD* elements with two *xis* phage integrase *anacy_2118* and *anacy_2114*. The lower of this scheme showed an alternative restoration of *primase P4* in this region by only removing *primase P4* element with a phage integrase *xis-anacy*_2120. **f)** The scheme showed the formation of intact *hupL* by removing the *hupL* element with phage integrase *xis*-*anacy*_1762. **g)** The scheme showed the formation of intact jointed *anacy_RS29550* & *RS_29775* hypothetical protein by removing the 39,998 bp element with a phage integrase *xis*-*anacy_6026*.

#### Primase P4 restoration editing in heterocysts and vegetative cells

*The primase P4* gene located between 5’-*nifD* and the middle part of *nifD* was interrupted in *A. cylindrica* by a *primase P4* DNA element (Fig. 3e). A 1,111 bp band was amplified with genomic DNA from vegetative cells grown in AA/8, AA/8N and heterocysts using primers ZR1748, ZR1749 (Fig. 3b). These three PCR products were sequenced and confirmed to be identical to the coding region of *primase P4* (GenBank accession #: MG594029). The qPCR data (Fig. 2) showed that 67.13±0.47% of *primase P4* was restored to be intact in heterocysts, while the restored *primase P4* gene in vegetative cells constituted only 0.03±0.004% (AA/8) and 0.03±0.003% (AA/8N) of total *primase P4* (interrupted and intact).

#### hupL restoration editing in heterocysts and vegetative cells

A 1,034 bp fragment (Fig. 3c) was amplified with genomic DNA from vegetative cells grown in AA/8, AA/8N and heterocysts with primers ZR1733, ZR1734 (Table S1). These three 1,034 bp PCR products were sequenced and confirmed to be the coding region of *hupL*. These results indicate that a 20,842 bp *hupL*-element, inserted within *hupL*, was precisely removed to restore an intact *hupL* (Fig. 3f & GenBank accession #: MG594030). Further qPCR data (Fig. 2) showed that 51.00±0.11% of *hupL* was edited to be intact in heterocysts, while the intact *hupL* in vegetative cells accounted for only 4.61±0.17% (AA/8) and 2.38±0.31% (AA/8N) of total *hupL* (interrupted and intact).

#### Hypothetical protein gene restoration editing in heterocysts and vegetative cells

A 744 bp fragment (Fig. 3d) was amplified with genomic DNA from vegetative cells grown in AA/8, AA/8N and heterocysts using primers ZR1744, ZR1745 (Fig. 2g & Table S1). These three 744bp PCR products were sequenced and confirmed that a 39,998 bp element in *pANACY.03* was precisely removed to form the 744 bp fragment. Thus, a hypothetical protein gene (1,539 bp) was restored by joining a 636 nt *anacy-RS29550* and a 903 nt *anacy_RS29775* (Fig. 3d &g and its GenBank accession #: MG594022). The qPCR data (Fig. 2) identified that the intact hypothetical protein gene accounted for 90.76±0.18% in heterocysts, while in vegetative cells accounted for 63.49±0.23% (AA/8) and 50.73±0.08% (AA/8N).

### Loss of 172 genes through the six DNA element deletions in heterocysts

The six DNA element deletions contained 156,752 bp and a total of 172 genes (Table S7). These deletions were identified by using *A. cylindrica* PCC 7122 as the reference genome. These 172 genes consist of 97 hypothetical protein genes or unknown genes; 19 phage-integrase genes or resolvase genes; 15 tRNA genes; 7 DNA-related genes involved in plasmid segregation (*Anacy_2158)*, DNA modification (*Anacy_6051)*, recombination (*Anacy_2126)*, and endonuclease activity (*Anacy_2171*, *2172*, *6054*, *6058*); 5 transposase genes; 5 ATPase genes; 5 transcription regulatory genes (*Anacy_1774*, *1777*, *1786*, *2168*, *2179*); 2 chromosome partitioning genes (*Anacy_2141*, *2142*); 2 prophage maintenance related genes (*Anacy_2147*, *2204*); 1 DNA replication-related gene (*Primase P4*); 1 photosystem gene (*Anacy_2162*); and 13 genes with other functions.

### Interrupted *nif* genes uniquely occur in heterocyst-forming N_2_-fixing cyanobacteria

Nearly all of the genes interrupted by DNA elements are known to be required for heterocyst-specific N_2_-fixation in 28 out of 38 heterocyst-forming cyanobacteria although a few of them have no known function in N_2_-fixation^26^. Since *nif* genes are highly conserved in all N_2_-fixing bacteria, why are the *nif* genes interrupted in most of the heterocyst-forming N_2_-fixing cyanobacteria, but not in non-heterocyst-forming cyanobacteria nor in other N_2_-fixing bacteria? We made an alignment for the three most commonly interrupted *nif* genes - *nifD* (Fig. S5abc), *nifH* (Fig. S6), and *hupL* (Fig. S7) - from the representatives of N_2_-fixing cyanobacterial species (multicellular and unicellular cyanobacteria) as well as non-photosynthetic N_2_-fixing bacteria (non-cyanobacteria) to identify if the direct repeat sequences (e.g., where the *nifD*-element inserted) were conserved in these orthologous *nif* genes.

For *nifD*, 25 direct repeat sequences (within the *nifD* insertion element) out of 32 interrupted *nifD* genes shared the conserved core sequence TACTCCG ^26^. Only one of the 32 interrupted *nifD* genes was interrupted by a serine integrase-containing DNA element, the rest were interrupted by a tyrosine integrase gene-containing DNA element ^26^. A 13-nucleotide sequence, including the core sequence (TACTCCG), in the *nifD* insertion element is targeted by tyrosine integrase. This nucleotide sequence was completely conserved in 16 heterocyst-forming cyanobacterial strains (Fig. S5a, boxed), indicating that these 16 *nifDs* are interrupted by a tyrosine integrase gene-containing DNA element ^26^. There were an additional 15 *nifD*s out of 40 uninterrupted *nifD*s (Fig. S5b) that also contain the conserved 13 nt sequence (*labeled in Fig. S5b), but have no interruption (Fig. S5b). These 15 *nifDs* were from eight heterocyst-forming cyanobacteria (five of *Fischerella* species and *Mastigocoleus testarum* BC008; *Calothrix desertica* PCC 7102; and *Nostoc azolla* 0708), three non-heterocyst filamentous N_2_-fixing cyanobacteria (*Oscillatoriales* JSC-12, two *Leptolyngbya* species) and four unicellular N_2_-fixing cyanobacteria (*Aphanocapsa* BDHKU 210001, *Chroococcidiopsis* PCC 7203, and two *Cyanothece* sp.) (Fig. S5b, *labeled). Only eight *nifDs* (Fig. S5c, *labeled) out of 71 uninterrupted *nifDs* from non-cyanobacteria also contain the absolutely conserved 13 nucleotide sequence TGGGATTACTCCG (Fig. S5a, boxed).

For *nifH*, seven *nifH* orthologs from three genera (Fig. S6a) were interrupted by a *nifH1* element containing a tyrosine integrase gene ^26^. All seven *nifHs* contained the absolutely conserved 29 nt sequence (Fig. S6a, boxed), including the direct repeat sequence CCGTGAAG where the *nifH* element inserted ^26^. Only five *nifH*-orthologs out of the 52 uninterrupted cyanobacterial *nifHs* (Fig. S6b,*labeled) contained the conserved 29 nt sequence (Fig. S6a, boxed). However, these five *nifHs* from heterocyst-forming *Ficherella* species and *Nostoc* species (Fig. S6b,*labeled) were not interrupted.

For *hupL*, ten *hupL*-orthologs from five genera (Fig. S7a) were interrupted by a *hupL* element containing a tyrosine integrase gene ^26^. All ten *hupL* genes contained the absolutely conserved 17 nt sequence (Fig. S7a, boxed) including the direct repeat sequence GCAGTTATATGG ^26^ where the *hupL*-element inserted (Fig. S7a, arrowed). Only five *hupL* orthologs out of the 30 uninterrupted cyanobacterial *hupL* (Fig. S7a,*labeled) contained the conserved 17 nt sequence (Fig. S7a, boxed). However, these five *hupL* from heterocyst-forming *Anabaena variabilis* and four *Cylindrospermopsis* species (Fig. S7b,*labeled) were not interrupted.

## Discussion

### *nif* gene editing may be accomplished in an early stage of heterocyst development

Heterocysts are specially differentiated, non-dividing cells with a unique function of solar-powered nitrogen fixation. It takes about 24 hours to develop a mature heterocyst from a vegetative cell of *A. cylindrica*. During heterocyst development, six large DNA elements (from 5,736 bp *nifH1*-element to 74,986 bp 5’-*nifD* element) that respectively interrupted five genes (*nifH1*, *nifD*, *hupL*, *primase P4*, and a hypothetical gene) in the genome of its progenitor vegetative cell are precisely deleted from the heterocyst genome of *A. cylindrica*. Thus, the five interrupted genes are restored to be functional in heterocysts. Our three levels of evidence: short reads mapping, contigs assembly and alignment to a reference genome, and quantitative PCR all confirmed these five genes’ restoration in heterocysts. However, our qPCR (Fig. 2) also detected that a small portion of genomic DNA from vegetative cells have these six DNA elements removed. For example, nearly 100% of interrupted *nifH*1 was edited to be intact in the genome of heterocysts (Fig. 2-*nifH1*), while only 11.02% (AA/8) and 4.86% (AA/8N) of interrupted *nifH*1 was edited to be intact in vegetative cells that are grown in the medium without combined nitrogen (AA/8) and with combined nitrogen (AA/8N). The percentage of the edited intact *nifH1* in vegetative cells had a positive correlation with the heterocyst frequencies (Fig. S1). That is, the higher heterocyst frequency culture had the higher *nifH1* edited in vegetative cells. Therefore, a small fraction of “vegetative cells” here might represent a stage of pro-heterocysts or an even earlier stage of heterocyst development. Although we cannot rule out the possibility of genome heterogeneity in vegetative cells due to the polyploidy nature of the genome and the multicellular morphology, our data supported that *nifH1* editing (*nifH*-element removal), like the other three genes (*nifD*, *hupL* and *primase P4*), was accomplished in a stage of pro-heterocyst or even earlier stage of heterocyst development.

Much research has demonstrated that nitrogen fixation in diazotrophic bacteria is tightly regulated at the transcriptional level ^30, 31, 32, 33, 34, 35, 36^. Here, we along with previous studies ^21, 22, 23, 37^ demonstrate that nitrogen fixation in heterocyst-forming cyanobacteria is also developmentally regulated at the genomic level. Our research shows that the interrupted *nif* genes in vegetative cells must be restored in heterocysts through genome editing during heterocyst development. The removal of the *nif*-elements (*nifD*, *fdxN)* from the heterocyst genome was found necessary for heterocyst N_2_-fixation, but not required for the differentiation of heterocysts in *Anabaena* sp. PCC 7120^23, 24^.

### The presence of different genomes in vegetative cells and heterocysts

It is generally believed that the distinct cell types in the same species should have the same genomes. However, recently, serval human cancer cells were reported to have very different genomes compared to the normal cells ^38, 39, 40, 41, 42^. Our comparative genomics between heterocysts and vegetative cells illustrated that there are at least 120 kb deletions in the heterocyst genome, including the six element deletions discussed above and another 33 potential deletions (Fig. S3). For heterocyst-forming cyanobacteria, these developmentally regulated genomic DNA deletions exclusively occurred in developing or mature heterocysts ^22, 37^, but have not yet been observed in vegetative cells. Similarly, during sporulation, regulated genomic DNA deletions (a 42.1-kb *spoVFB* element and a 48-kb *sigK* element) occurred exclusively in mother cells of *Bacillus weihenstephanensis* KBAB4 and *B. subtilis*, but were not seen in their forespores ^21, 43, 44^. Interestingly, both the heterocysts and the mother cells are terminally differentiated, non-dividing cells, unable to produce a next generation and eventually die. Therefore, the genomic DNA continuity is preserved in vegetative cells in heterocyst-forming cyanobacteria and in spores of spore-forming *Bacilli* ^43, 44^.

Among the six DNA elements (*nifH1*, *nifD*, *hupL*, *primase P4* and a hypothetical protein gene), at least one phage integrase gene was associated with each DNA element. Additionally, some prophage related genes (*Anacy_2147*, *2204*) were found within the DNA elements. The six DNA elements may have originated from a prophage or prophage remnants. Thus, the *nif* genes were interrupted by insertion of prophage DNA, causing oxygen (O_2_)-sensitive nitrogen fixation to be silenced in O_2_-producing vegetative cells, and restored in terminally differentiated heterocysts by removing these elements within the *nif* genes from the heterocyst genome. At the same time, some functions such as photosynthesis, DNA replication, etc. could be deactivated in heterocysts due to a loss of the 172 genes, with 97 of these genes having unknown function (Table S7). Clearly *nif* gene restoration (removal of *nif* elements) in *Anabaena* sp. is a developmentally regulated event, but the signal triggering this event remains unknown. Given the DNA elements’ prophage identity and the deletions of DNA elements not required for heterocyst differentiation in cyanobacteria ^23, 24, 45^, we hypothesize that such a signaling may come from an integrated collaborative decision between the cryptic lysogenic phages and developing heterocysts.

### *nif* genes interrupted by insertion of prophage DNA occurs uniquely in heterocyst-forming N_2_-fixing cyanobacteria

Why does the interruption of *nif* genes by a prophage DNA uniquely occur in heterocyst-forming cyanobacteria? A better interpretation of these insertions is that they are cryptic lysogenic phages that have found a non-lethal, highly conserved target site within the *nif* genes, at the cost of excising in terminally differentiated cells, the heterocysts. If this is true, a completed heterocyst genome compared to its vegetative cells genome may help us discover more lysogenic phages.

Using *nifD* gene as an example, we made a sequence alignment for 128 *nifD* genes from the representatives of three groups of N_2_-fixing bacteria (heterocyst-forming cyanobacteria, non-heterocyst-forming cyanobacteria, non-cyanobacteria). This sequence alignment revealed that only 39 *nifD* genes contained the conserved 13 nt TGGGATTACTCCG (Fig. S5a) found in the 16 interrupted *nifD* genes. Twenty-four out of the 39 were from heterocyst-forming cyanobacteria, with 16 interrupted by phage DNA and 8 uninterrupted (*labeled in Fig. S5b) from 5 *Fischerella* species (PCC 73103, PCC 605, PCC 7414, PCC 7512, and JSC-11), *Mastigocoleus testarum* BC008; *Calothrix desertica* PCC 7102 and *Nostoc azolla* 0708. The remaining 15 were from non-heterocyst forming cyanobacteria or non-cyanobacteria. None of these 15 *nifD* genes were interrupted by phage DNA.

For the 8 uninterrupted *nifD* genes from heterocyst-forming cyanobacteria, we tried to understand why they were not interrupted. First, none of the genomes of the 5 *Fischerella* species had any DNA element insertions. Neither did the genome of *Mastigocoleus testarum* BC 008 ^26^. Second, *Nostoc azolla* 0708 is a heterocyst-forming cyanobacterium that forms an endosymbiotic relationship with a plant, the water-fern *Azolla filiculoides Lam* ^46^; thus the cyanophage may not have access to the inside of a plant cell. Currently, we are unable to hypothesize why *Calothrix desertica* PCC 7102 contains uninterrupted *nif* genes. Although its genome contains eight DNA element insertions found in other locations ^26^, none of them inserted within *nifD*.

For the 15 *nifD* from non-heterocyst forming cyanobacteria and other N_2_-fixing bacteria, none were interrupted by phage DNA. We speculate that the conserved 13 nt sequence is not all that is required for integration of the *nifD* element. Several other factors, including but not limited to an integration host factor, integrase, and excisionase, are required. Evolutionary selection would also play an important role in the DNA element integration. For example, if a *nif* gene is interrupted by a prophage DNA in non-heterocystous N_2_-fixing cyanobacteria or other N_2_-fixing bacteria, it would place them at a disadvantage to the bacteria that retain an intact nitrogenase gene. This would also present a selective pressure in heterocyst-forming cyanobacteria, but due to their cooperative multicellular morphology, the selective pressure would not be as severe as it would be for a non-heterocystous cyanobacteria or non-cyanobacteria. In other words, the prophage DNA might once have inserted within these 15 *nifD*s, but natural selection pressure on non-heterocystous cyanobacteria made these insertion mutants unfit (i.e., forced them to become extinct) in a combined-nitrogen-free environment.

## Author Contributions

YQ, LG, and RZ designed the work. YQ, LG, ST, and JG performed the experiments. YQ, LG, JS, TD, JGH and RZ analyzed the genomic data and drafted the manuscript. YQ, LG, JS, JGH, JG and RZ revised the manuscript and are responsible for final approval of the version to be published. All authors agree to be accountable for the content of the work.

## Acknowledgements

This work was partially supported by USDA-NIFA GRANT Heterocyst Transcriptomics (to R. Z.), and by the South Dakota Agricultural Experiment Station. The authors would like to acknowledge use of the South Dakota State University Functional Genomics Core Facility supported in part by NSF/EPSCoR Grant No. 0091948 and by the State of South Dakota.

## References

1. Wolk CP. Heterocyst formation. Annu Rev Genet 30, 59–78 (1996).

2. Golden JW, Yoon HS. Heterocyst development in Anabaena. Curr Opin Microbiol 6, 557–563 (2003).

3. Meeks JC, Wycoff KL, Chapman JS, Enderlin CS. Regulation of expression of nitrate and dinitrogen assimilation by anabaena species. Appl Environ Microbiol 45, 1351–1359 (1983).

4. Murry MA, Wolk CP. Evidence that the barrier to the penetration of oxygen into heterocysts depends upon two layers of the cell envelope. Archives of Microbiology 151, 469–474 (1989).

5. Zhou RB, Wolk CP. A two-component system mediates developmental regulation of biosynthesis of a heterocyst polysaccharide. Journal of Biological Chemistry 278, 19939–19946 (2003).

6. Walsby AE. Cyanobacterial heterocysts: terminal pores proposed as sites of gas exchange. Trends Microbiol 15, 340–349 (2007).

7. Adams DG, Carr NG. Control of heterocyst development in the cyanobacterium Anabaena cylindrica. Microbiology 135, 839–849 (1989).

8. Buikema WJ, Haselkorn R. Characterization of a gene controlling heterocyst differentiation in the cyanobacterium Anabaena 7120. Genes Dev 5, 321–330 (1991).

9. Zhou R, et al. Evidence that HetR protein is an unusual serine-type protease. Proc Natl Acad Sci U S A 95, 4959–4963 (1998).

10. Huang X, Dong Y, Zhao J. HetR homodimer is a DNA-binding protein required for heterocyst differentiation, and the DNA-binding activity is inhibited by PatS. Proc Natl Acad Sci U S A 101, 4848–4853 (2004).

11. Ehira S, Ohmori M. NrrA, a nitrogen-regulated response regulator protein, controls glycogen catabolism in the nitrogen-fixing cyanobacterium Anabaena sp. strain PCC 7120. J Biol Chem 286, 38109–38114 (2011).

12. Hu Y, et al. Structures of Anabaena calcium-binding protein CcbP: insights into Ca2+ signaling during heterocyst differentiation. J Biol Chem 286, 12381–12388 (2011).

13. Higa KC, et al. The RGSGR amino acid motif of the intercellular signalling protein, HetN, is required for patterning of heterocysts in Anabaena sp. strain PCC 7120. Mol Microbiol 83, 682–693 (2012).

14. Risser DD, Callahan SM. HetF and PatA control levels of HetR in Anabaena sp. strain PCC 7120. Journal of bacteriology 190, 7645–7654 (2008).

15. Risser DD, Wong FC, Meeks JC. Biased inheritance of the protein PatN frees vegetative cells to initiate patterned heterocyst differentiation. Proc Natl Acad Sci U S A 109, 15342–15347 (2012).

16. Meeks JC, Campbell EL, Summers ML, Wong FC. Cellular differentiation in the cyanobacterium Nostoc punctiforme. Arch Microbiol 178, 395–403 (2002).

17. Zhang W, et al. A gene cluster that regulates both heterocyst differentiation and pattern formation in Anabaena sp. strain PCC 7120. Mol Microbiol 66, 1429–1443 (2007).

18. Hu HX, et al. Structural insights into HetR-PatS interaction involved in cyanobacterial pattern formation. Sci Rep 5, 16470 (2015).

19. Yoon HS, Golden JW. Heterocyst pattern formation controlled by a diffusible peptide. Science 282, 935–938 (1998).

20. Videau P, et al. The heterocyst regulatory protein HetP and its homologs modulate heterocyst commitment in Anabaena sp. strain PCC 7120. Proc Natl Acad Sci U S A, (2016).

21. Haselkorn R. Developmentally regulated gene rearrangements in prokaryotes. Annu Rev Genet 26, 113–130 (1992).

22. Golden JW, Robinson SJ, Haselkorn R. Rearrangement of nitrogen fixation genes during heterocyst differentiation in the cyanobacterium Anabaena. Nature 314, 419–423 (1985).

23. Golden JW, Wiest DR. Genome rearrangement and nitrogen fixation in Anabaena blocked by inactivation of xisA gene. Science 242, 1421–1423 (1988).

24. Carrasco CD, Ramaswamy KS, Ramasubramanian TS, Golden JW. Anabaena xisF gene encodes a developmentally regulated site-specific recombinase. Genes Dev 8, 74–83 (1994).

25. Carrasco CD, Holliday SD, Hansel A, Lindblad P, Golden JW. Heterocyst-specific excision of the Anabaena sp. strain PCC 7120 hupL element requires xisC. Journal of bacteriology 187, 6031–6038 (2005).

26. Hilton JA, Meeks JC, Zehr JP. Surveying DNA Elements within Functional Genes of Heterocyst-Forming Cyanobacteria. PLoS One 11, e0156034 (2016).

27. Hu NT, Thiel T, Giddings TH, Jr., Wolk CP. New Anabaena and Nostoc cyanophages from sewage settling ponds. Virology 114, 236–246 (1981).

28. Lawrence M, et al. Software for computing and annotating genomic ranges. PLoS Comput Biol 9, e1003118 (2013).

29. Kurtz S, et al. Versatile and open software for comparing large genomes. Genome Biol 5, R12 (2004).

30. Dixon R, Kahn D. Genetic regulation of biological nitrogen fixation. Nature Reviews Microbiology 2, 621 (2004).

31. Shi HW, et al. Genome-wide transcriptome profiling of nitrogen fixation in Paenibacillus sp. WLY78. BMC Microbiol 16, 25 (2016).

32. Flaherty BL, Van Nieuwerburgh F, Head SR, Golden JW. Directional RNA deep sequencing sheds new light on the transcriptional response of Anabaena sp. strain PCC 7120 to combined-nitrogen deprivation. BMC Genomics 12, 332 (2011).

33. Vernon SA, Pratte BS, Thiel T. Role of the nifB1 and nifB2 Promoters in Cell-Type-Specific Expression of Two Mo Nitrogenases in the Cyanobacterium Anabaena variabilis ATCC 29413. J Bacteriol 199, (2017).

34. Tsujimoto R, Kamiya N, Fujita Y. Transcriptional regulators ChlR and CnfR are essential for diazotrophic growth in nonheterocystous cyanobacteria. Proc Natl Acad Sci U S A 111, 6762–6767 (2014).

35. Tsujimoto R, Kamiya N, Fujita Y. Identification of a cis-acting element in nitrogen fixation genes recognized by CnfR in the nonheterocystous nitrogen-fixing cyanobacterium Leptolyngbya boryana. Mol Microbiol 101, 411–424 (2016).

36. Poza-Carrion C, Jimenez-Vicente E, Navarro-Rodriguez M, Echavarri-Erasun C, Rubio LM. Kinetics of Nif gene expression in a nitrogen-fixing bacterium. J Bacteriol 196, 595–603 (2014).

37. Carrasco CD, Buettner JA, Golden JW. Programmed DNA rearrangement of a cyanobacterial hupL gene in heterocysts. Proc Natl Acad Sci U S A 92, 791–795 (1995).

38. Barbieri CE, Rubin MA. Genomic rearrangements in prostate cancer. Curr Opin Urol 25, 71–76 (2015).

39. Network CGAR. Comprehensive genomic characterization of squamous cell lung cancers. Nature 489, 519–525 (2012).

40. Periwal V, Scaria V. Insights into structural variations and genome rearrangements in prokaryotic genomes. Bioinformatics 31, 1–9 (2015).

41. Smith JJ, Baker C, Eichler EE, Amemiya CT. Genetic consequences of programmed genome rearrangement. Curr Biol 22, 1524–1529 (2012).

42. Stratton MR, Campbell PJ, Futreal PA. The cancer genome. Nature 458, 719–724 (2009).

43. Abe K, Yoshinari A, Aoyagi T, Hirota Y, Iwamoto K, Sato T. Regulated DNA rearrangement during sporulation in Bacillus weihenstephanensis KBAB4. Mol Microbiol 90, 415–427 (2013).

44. Stragier P, Kunkel B, Kroos L, Losick R. Chromosomal rearrangement generating a composite gene for a developmental transcription factor. Science 243, 507–512 (1989).

45. Kunkel B, Losick R, Stragier P. The Bacillus subtilis gene for the development transcription factor sigma K is generated by excision of a dispensable DNA element containing a sporulation recombinase gene. Genes Dev 4, 525–535 (1990).

46. Ran L, et al. Genome erosion in a nitrogen-fixing vertically transmitted endosymbiotic multicellular cyanobacterium. PLoS One 5, e11486 (2010).

